# Butyrate-producing bacteria as probiotic supplement: beneficial effects on metabolism and modulation of behavior in an obesity mouse model

**DOI:** 10.1101/2023.11.02.564919

**Authors:** Alba M. Garcia-Serrano, Cecilia Skoug, Ulrika Axling, Emma Rosmariini Korhonen, Cristina Teixeira, Irini Lazou Ahrén, Indrani Mukhopadhya, Nikoleta Boteva, Jennifer Martin, Karen Scott, Silvia Gratz, Karin G. Stenkula, Cecilia Holm, João M.N. Duarte

## Abstract

Obesity is a risk factor for cardio-metabolic and neurological disease. The contribution of gut microbiota to derangements of the gut-brain axis in the context of obesity has been acknowledged, particularly through physiology modulation by short-chain fatty acids (SCFAs). Thus, probiotic interventions and administration of SCFAs have been employed with the purpose of alleviating symptoms in both metabolic and neurological disease. We have tested the effects of four butyrate-producing bacteria from the Lachnospiraceae family on the development of metabolic syndrome and behavioural alterations in a mouse model of diet-induced obesity. Male mice were fed either a high-fat diet (HFD) or an ingredient-matched control diet for 2 months, and bacteria cultures or culture medium were given by gavage to HFD-fed mice every second day. Assessment of the mice through a battery of metabolic and behaviour tests revealed that one of the administered bacteria affords some degree of protection against the development of obesity and its complications.

## Introduction

Overweight and obesity pose a major risk for serious comorbidities such as cardiovascular disease, hypertension and stroke, metabolic syndrome and diabetes (Schlesinger *et al*., 2021). These diseases, in turn, all affect the brain and are a risk for dementia (Anstey *et al*., 2011; Albanese *et al*., 2017). The dramatic increase in obesity rates is mostly associated with high intake of food products rich in saturated fat and sugar, and with low physical activity and sedentary behaviour. Thus, it is imperative to design strategies for healthy lifestyle choices, as well as to develop functional food products that prevent obesity and associated comorbidities.

The gut harbours symbiotic bacteria, archaea and eukarya (called microbiota) that modulate human physiology, and have been strikingly associated with pathological processes including the development of obesity, diabetes and related comorbidities (Brunkwall & Orho-Melander, 2017; Amabebe *et al*., 2020), and with neurodegenerative disorders (Morais *et al*., 2020). Biochemical and hormonal signalling occurs between microbiota in the gastrointestinal tract and the central nervous system, the so-called microbiome-gut-brain axis. Moreover, there is a crosstalk between intestinal microbiota and the hypothalamic-pituitary-adrenal (HPA) axis that controls energy homeostasis and stress responses (Carabotti *et al*., 2015; Morais *et al*., 2020). The gut-brain connection is facilitated by the vagus nerve and enteric nervous system, as well as endocrine, immune and metabolic pathways, but many of its molecular mediators remain to be identified.

Efforts have been conducted to elucidate how microbiome alterations contribute to metabolic and neurological disorders, and specific probiotic supplementations might have potential for disease prevention and management (Amabebe et al., 2020). In recent years, special interest has focused on health benefits from the so-called next generation probiotics (O’Toole et al., 2017). In contrast to traditional probiotic bacteria, usually lactobacilli or bifidobacteria, next generation probiotics represent bacteria from other genera such as *Bacteroides*, *Akkermansia* and *Eubacterium* that grow anaerobically and have been isolated relatively recently.

Gut bacteria produce many metabolites from dietary sources, including short-chain fatty acids, such as butyrate, which are functional modulators and fuel sources for intestinal epithelial cells, and moreover impact human health by acting on host immunity, metabolic regulation, and brain function (Martin-Gallausiaux *et al*., 2021; van der Hee & Wells, 2021). Butyrate is generally regarded as a metabolite that can alleviate obesity and associated metabolic derangements, although some studies dispute this (Liu *et al*., 2018). In this study, we aimed at testing the efficacy of four anaerobic butyrate-producing bacterial strains from the Lachnospiraceae family in promoting metabolic health, and improving brain function. For that, we exposed mice to a high-fat diet (HFD), which is a well-characterized model of obesity with metabolic syndrome (Soares *et al*., 2018) and memory impairment (Garcia-Serrano, Vieira *et al*., 2022), and treated them with any of these four bacterial strains for two months. Afterwards, we evaluated metabolic health, and brain function and behaviour phenotypes such as memory, anxiety and depression.

## Materials and Methods

### Bacterial suspension

Four anaerobic bacteria from the Lachnospiraceae family (B1, B2, B3 and B4) had been previously isolated from faecal or biopsy samples of healthy volunteers (Mukhopadhya *et al*., 2022), and were selected for this study based on their ability to produce >10 mmol/L butyric acid in YCFAGSC media. Each bacterium was grown in Hungate tubes containing 7 mL anaerobic YCFAGSC medium (Duncan *et al*., 2002), for 16 hours at 37°C up to their maximum optical density at 600 nm with an average of 2.2±0.5, 1.0±0.3, 1.7±0.2 and 2.7±0.3 for B1, B2, B3 and B4, respectively.

### Animals

Mouse experiments were performed according to EU Directive 2010/63/EU under approval by the Malmö/Lund Committee for Animal Experiment Ethics (permit number 5123/2021), and are reported following the ARRIVE guidelines (Animal Research: Reporting In Vivo Experiments, NC3Rs initiative, UK). Male C57BL/6J mice (8-weeks old) purchased from Taconic Biosciences (Köln, Germany) were housed in groups of 4-5 on a 12-hour light-dark cycle with lights on at 07:00, room temperature of 21-23°C and humidity at 55-60%. Cages were furnished with a climbing structure, cylinder, wood toys and nesting material. Mice were habituated to the facility for a week upon arrival. Cages were randomly assigned to experimental groups using the RANDBETWEEN function in Excel (Microsoft, Redmond, WA-USA).

Dietary intervention and bacteria supplementation started at 9 weeks of age and continued for 2 months (study design in Figure 1A). Food and water were provided *ad libitum*. Diets were acquired from Research Diets (New Brunswick, NJ-USA): a lard-based diet with 60% kcal of fat (HFD) (D12492) and a control diet (CD) containing 10% kcal of fat (D12450J), with total energy of 5.21 and 3.82 kcal/g, respectively. Gross caloric intake was measured as the difference in weight between the food put into the cage and that remaining at the end of the week. Researchers knew the group allocation during experiments, but not during sample processing and data analysis. This allowed the sequence of mouse handling to be shuffled, thus minimizing potential confounders.

**Figure 1.**
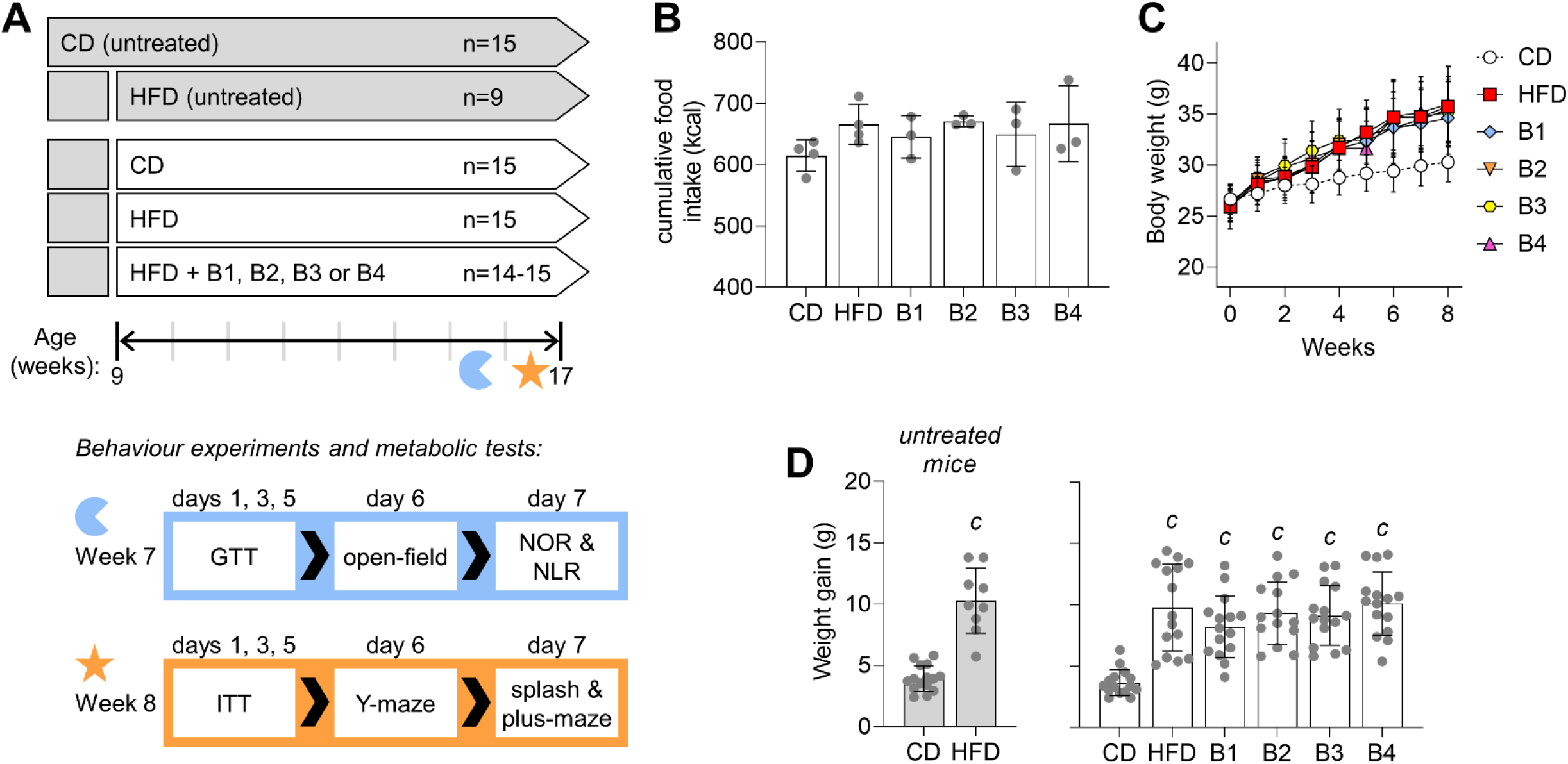
Study design, and timing of experiments during the last two weeks of dietary intervention and treatment. Two groups of mice were kept untreated, that is, have not been gavaged throughout the study. Six groups of mice were gavaged every second day with either culture medium or bacterial suspension during whole period of exposure to HFD. Mice went through GTT and ITT on days of gavage (A). Cumulative intake of calories as measured from cage-average food consumption (n=3-4 cages, B). Body weight of mice receiving vehicle or probiotic preparations via oral gavage (C), and weight gain from baseline for treated and untreated mice (D). Data are shown as mean ± SD, with dots representing each mouse (n=9-15, as indicated in A). *^c^*P<0.001 for comparison to CD based on Student t-tests, or on Fischer’s LSD post-hoc testing following a significant ANOVA effect.

Mice received an oral gavage of 0.2 mL of bacterial suspension or culture medium (vehicle) between 8:00 and 11:00 every other day during the HFD-diet intervention. A parallel cohort of mice fed CD or HFD was left untreated as control for gavage and handling.

### Behavior testing

Mice were allowed to acclimatize to the testing room for 1 hour before each experiment, and tests were performed from 9:00 to 18:00, with room light adjusted to achieve an illuminance of 15 lx in the test apparatus, as previously described (Garcia-Serrano, Vieira *et al*., 2022). Experiments were recorded by an infrared camera into the ANY-maze software (6.0.1, Stoelting).

Open field test and object recognition tasks for testing novel object recognition (NOR) or novel location recognition (NLR) were conducted as detailed elsewhere (Garcia-Serrano, Vieira *et al*., 2022). Mice were first habituated to the empty open-field arena for 8 minutes. Arena exploration was tracked and analysed with ANY-maze 6.0 (Dublin, Ireland). Total walk distance, number of crossings between arena quadrants and immobility time, as well as exploration of the arena centre at 6 cm from the walls were tracked. Thereafter, NLR was assessed by placing the mice in the arena with two identical objects, and allowing them to explore these for 5 minutes (familiarization phase). Mice were then removed from the arena for 1 hour (retention phase), and reintroduced for 5 minutes but with one object relocated to a different quadrant in the arena (recognition phase). For NOR, two new identical objects were used in the familiarization phase, and one of them was replaced by a novel object during the recognition phase. Time exploring each object was measured.

Spontaneous alternations were observed in a Y-maze as surrogate of working memory performance (Duarte *et al*., 2012). The Y-maze arms were 30 cm x 15 cm x 5 cm (length x height x wide), and converged to the center in 120° angles. Mice were placed in one arm of the maze and allowed to freely explore for 5 minutes. A complete spontaneous alternation was defined as a successive entrance to each different arm, and expressed relative to the total possible alternations in the respective test. The total number of entries was analysed to access locomotor activity and exploratory behaviour.

The Elevated Plus Maze was used to assess anxiety as previously described (Skoug *et al*., 2022). Each maze arm was 35 cm x 5 cm, and closed arms had 15 cm walls, at a 60 cm height from the floor. The mouse was placed in the maze center facing an open arm, and was allowed to freely explore the maze for 5 minutes. Number of entries and time spent in each arm were analysed.

Self-care behaviour was analysed with the sucrose splash test, as a means of inferring on a depression-like phenotype (de Paula *et al*., 2020). Mice were splashed with a 10% (w/v) sucrose solution on their dorsal coat, and latency to grooming, and grooming duration were recorded during 5 minutes.

### Glucose tolerance test (GTT) and Insulin tolerance test (ITT)

For GTT, food was removed for 6 hours starting at 08:00. After, fasting glycemia was measured from the tail tip blood with a Breeze glucometer (Bayer, Zürich, Switzerland), blood from the vena saphena was collected into a heparinized tube for determining plasma insulin by ELISA (#10-1247-10, Mercodia, Uppsala, Sweden), and a GTT with 2 g/kg glucose given intra-peritoneally (i.p.) was carried out as previously described (Garcia-Serrano, Mohr *et al*., 2022). For ITT, we measured fed glycemia and collected a sample for insulin determination as above. Then, mice were injected i.p. with 2 U/kg of an insulin analogue (Actrapid 100 U/mL, Novo Nordisk Scandinavia, Malmö, Sweden) from a 0.125 U/mL solution in saline. In both GTT and ITT, tail-tip blood glucose levels were determined at 15, 30, 60, 90 and 120 minutes post-injection.

### Dual-energy X-ray absorptiometry (DEXA)

Dual-energy x-ray absorptiometry (DEXA) was performed under brief ∼2% isoflurane anesthesia in air, with mice positioned ventrally in a Lunar PIXImus2 scanner (GE Healthcare, Danderyd, Sweden). The mouse head was partially outside the field of view, and thus it was completely excluded from further analysis by placing an exclusion region of interest, according to the manufacturer’s instructions. Parameters estimated were total bone mineral density and mass, bone-free lean tissue mass, and fat mass.

### Gut barrier and blood-brain barrier (BBB) permeability

Fluorescein isothiocyanate (FITC)-dextran of 4,000 Da (#46944, Sigma-Aldrich) and Rhodamine B-Dextran of 70,000 Da (#D1841, Invitrogen, ThermoFisher) were used for determining permeability of the gut barrier and BBB, respectively. Mice received 100 µL i.p. of 40 g/L Rhodomine B-dextran in phosphate-buffered saline (PBS; in mmol/L: 137 NaCl, 2.7 KCl, 1.5 KH_2_PO_4_, 8.1 Na_2_HPO_4_, pH 7.4), followed by 100 µL gavage of 100 g/L FITC-dextran in PBS by oral gavage. After 60 minutes, mice were anesthetized with 2% isofluorane, and blood samples were collected from the heart and the portal vein into heparinized, light-protected tubes, and plasma was frozen. After transcardiac perfusion with cold PBS, brain structures were dissected in the penumbra and immediately frozen in N_2_ (l). Brain tissue samples were homogenized in 150 µL of PBS, and centrifuged at 18,000 g for 20 minutes. Supernatants and plasma samples diluted 6-fold in PBS were used to measure Rhodamine B and FITC fluorescence in 96-well opaque plates using a Fluostar Ω microplate reader (BMG Labtech, Ortenberg, Germany).

### Liver triglycerides

Liver triglyceride concentration was determined as described previously (Fryklund *et al*., 2022). In short, frozen liver samples (∼100 mg) were homogenized using a Dounce homogenizer in 5% (w/v) NP-40 lysis buffer. The homogenized samples were heated at 90°C for 3 minutes, cooled down for 15 minutes at room temperature, reheated at 90°C for 3 minutes, and then centrifuged at 10,000 *g* for 2 minutes at 20°C. The concentration of triglycerides in supernatants was determined using a kit from ThermoFisher (#TR22421) and calibration standards prepared from normal control serum (#TR40001).

### Caco-2 cell culture

Caco-2 cells were obtained from the European Collection of Authenticated Cell Cultures (ECACC 86010202) through Sigma-Aldrich (Stockholm, Sweden). Cells were routinely maintained in 75-cm^2^ flasks in Dulbecco’s modified Eagle’s medium (DMEM) with 4.5 g/L D-glucose, L-glutamine, sodium pyruvate (#6429, Sigma-Aldrich) supplemented with 1% (v/v) non-essential amino acids (NEAA) (Gibco, #11140-035), 1% (v/v) glutagro (Corning, #25-015-Cl), penicillin 10,000 U/mL and streptomycin 10,000 U/mL (Cytiva, #SV30010), and 10% (v/v) heat-inactivated fetal bovine serum (Sigma-Aldrich, #7524) in a humidified atmosphere of 5% CO_2_ at 37°C. Cells were subcultured using 0.25% Trypsin-EDTA (Gibco, #25200-056) at 80-90% confluency using a split of 1:5 to 1:10. For experiments, cells were seeded in Corning Transwell 12-well inserts with a polyester membrane (0.4 μm pore size, 1.12 cm^2^ surface area) at a density of 5x10^5^ cells/well. Medium was changed every third day and experiments were performed 21 days after seeding.

### Caco-2 cell treatments and transepithelial electrical resistance (TEER) measurement

B1 was grown as described above to an optical density of 2.4 (absorbance at 600 nm). To prepare the different bacterial fractions, bacterial cultures were centrifugated at 3200 x *g* for 6 minutes at room temperature. The supernatant was collected, sterile filtered through a polyethersulfone membrane with 0.2 μm pore size, and defined as the supernatant fraction. The bacterial pellet was washed twice and resuspended to its original volume in serum-free cell culture medium. Heat-inactivation was performed at 70°C for 3 minutes. On the day of experiment, medium was replaced with serum-free medium and after equilibration to room temperature, TEER was measured using a Millicell ERS-2 epithelial volt-ohm meter (Millipore). TEER was recorded three times in each well, in three different positions, and an average was calculated. After removal of 100 μL of medium from the apical compartments, bacterial fractions were added. Serum-free cell culture medium and bacterial broth, respectively, were used as controls. Following 24 hours of incubation in a humidified atmosphere of 5% CO_2_ at 37°C, plates were equilibrated to room temperature and TEER was measured. To challenge the cell barrier, TNF-α (Sigma-Aldrich, #H8916) at a concentration of 2 ng/mL was added alone or together with 30 ng/mL IFN-γ (Sigma-Aldrich, #I7001) and 3 ng/mL IL-1β (Sigma-Aldrich, #H6291), and the incubation was extended for another 24 to 96 hours, after which TEER was measured again. In each set of experiments inserts without cells were included to allow for correction of the TEER value for background resistance. The resistance values (Ω cm^2^) of the cell monolayer was obtained by subtracting the average TEER measurements of the blank inserts and multiplying by the surface area of the insert. All cell monolayers presented TEER values above 500 Ω cm^2^ at the start of the experiment and typically reached maximum values above 3000 Ω cm^2^.

### SCFA analysis

Concentrations of short chain and branched chain fatty acids (SCFA and BCFA respectively) in cell culture supernatants were determined by gas chromatography as previously described (Mukhopadhya *et al*., 2022, and references therein). Briefly, after derivatisation using *N*-tert-butyldimethylsilyl-*N*-methyltrifluoroacetamide, samples were analysed using a Hewlett Packard (Palo Alto, CA-USA) gas chromatograph fitted with a fused silica capillary column using helium as the carrier gas. Concentrations were calculated from the relative response factor relative to the internal standard 2-ethylbutyrate and a mixture of external standards (6 SCFAs in water).

### Statistical analyses

Results are presented as mean±SD unless otherwise stated, and were analyzed with Prism 9.0.2 (GraphPad, San Diego, CA-US). Kolmogorov-Smirnov and Shapiro-Wilk tests were used for inferring on normality. In the absence of normality deviations, results were analyzed using unpaired, 2-tailed Students t-test or ANOVA followed by independent comparisons with the Fisher’s least significant difference (LSD) test. Kruskal-Wallis followed by Dunn’s post-tests were used for non-Gaussian-distributed data. Significance was accepted for P<0.05.

A partial least-squares (PLS) regression with 5 components was applied on z-scores of measured parameters for CD and HFD groups using MATLAB 2019a (MathWorks, Natick, MA-USA). For that, each sample was assigned a dummy variable as a reference value (0 = CD and 1 = HFD). The variable importance in projection (VIP) was calculated for each parameter. The obtained PLS regression model was used to determine the component space for mice in groups treated with the bacteria suspension.

## Results

### Metabolic phenotype

Energy intake calculated from food consumption across the 8 weeks of treatment was largely similar between the experimental groups (ANOVA F=1.104, P=0.40; figure 1B). Body weight gain was larger for mice under HFD than CD (figure 1C), with total 8-week weight gain being 2-fold larger in HFD-fed mice than controls, and was not affected by the bacterial suspension supplementations (ANOVA F=13.26, P<0.001; figure 1D). Similar weight gain difference was observed for untreated mice, suggesting that oral gavage every other day was devoid of any effect on food consumption.

As expected, HFD-induced obesity was accompanied by increased fasting glycemia (ANOVA F=5.751, P<0.001; figure 2A), and increased fasting insulin levels in plasma (Kruskal-Wallis P<0.001; figure 2B). Glucose clearance after an oral glucose load was impaired upon HFD consumption (figure 2C), as evidenced by the larger area under the curve of the glucose excursion curves (ANOVA F=7.013, P<0.001; figure 2D). Compared to CD,HFD-fed mice also showed impaired glycemic response to injection of insulin (figure 2E), namely with typically higher plasma glucose after 30 minutes in HFD-fed mice under any treatment *versus* controls (ANOVA F=6.399, P<0.001; figure 2F). Interestingly, treatment with B1, but none of the other bacteria tested, lowered HFD-induced increased glycemia, ameliorated glucose clearance, and improved insulin sensitivity (figure 2).

**Figure 2.**
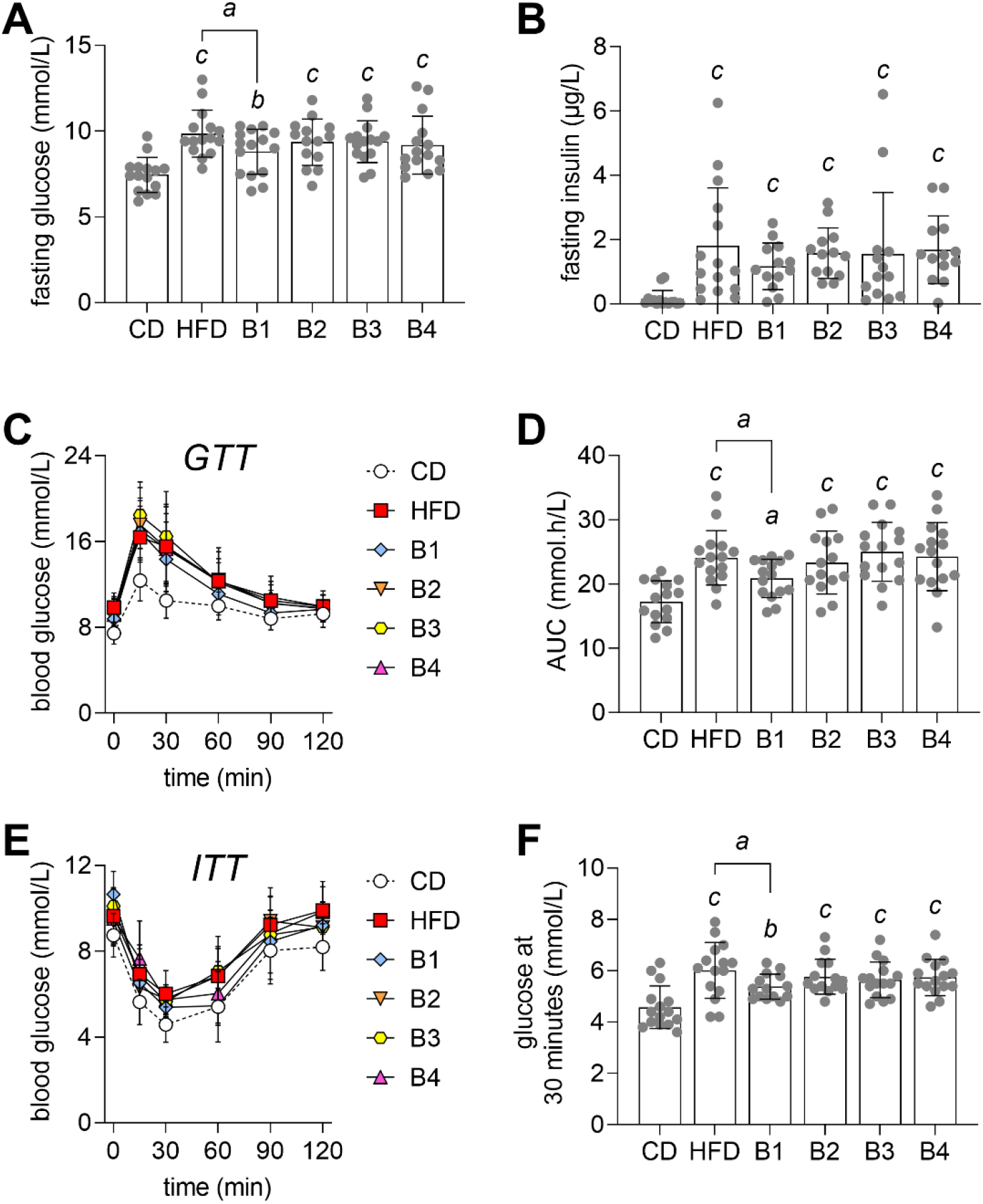
Fasting concentrations of glucose in blood (A) and insulin in plasma (B). Glucose clearance in a GTT (C), and respective area under the curve (AUC) of the GTT (D). Blood glucose levels during the insulin tolerance test (E), and glycemia at 30 minutes after insulin injection (F). Data are shown as mean ± SD, with dots representing each mouse (n=14-15). In A, D and F, letters over bars indicate significant differences (*^a^*P<0.05, *^b^*P<0.01, *^c^*P<0.001) for comparison to CD or as indicated, based on Fischer’s LSD post-hoc testing following a significant ANOVA effect. For B, insulin concentrations showed non-Gaussian distribution and were analysed with the Kruskal-Wallis test followed by Dunn’s multiple comparisons.

HFD exposure resulted in increased adiposity. Compared to controls, HFD-fed mice showed increased epididymal fat accumulation (ANOVA F=13.29, P<0.001; figure 3A) and whole-body fat content (ANOVA F=20.16, P<0.001; figure 3B), without changes in total lean mass (ANOVA F=0.447, P=0.81; figure 3C). DEXA scans showed similar bone mineral content (ANOVA F=0.69, P=0.64; figure 3D) and density (ANOVA F=1.90, P=0.10; figure 3E) across the experimental groups. Similar spleen size independently of treatments suggest minimal inflammation in HFD (ANOVA F=1.13, P=0.27; figure 3F). HFD had negligible effects on liver weight (ANOVA F=1.90, P=0.10; figure 3G), although hepatic triglyceride content was increased in all HFD-fed groups (ANOVA F=2.70, P=0.026; figure 3H). In line with findings on glucose tolerance, treatment with B1 reduced the HFD-induced increase in whole-body fat content (figure 3B).

**Figure 3.**
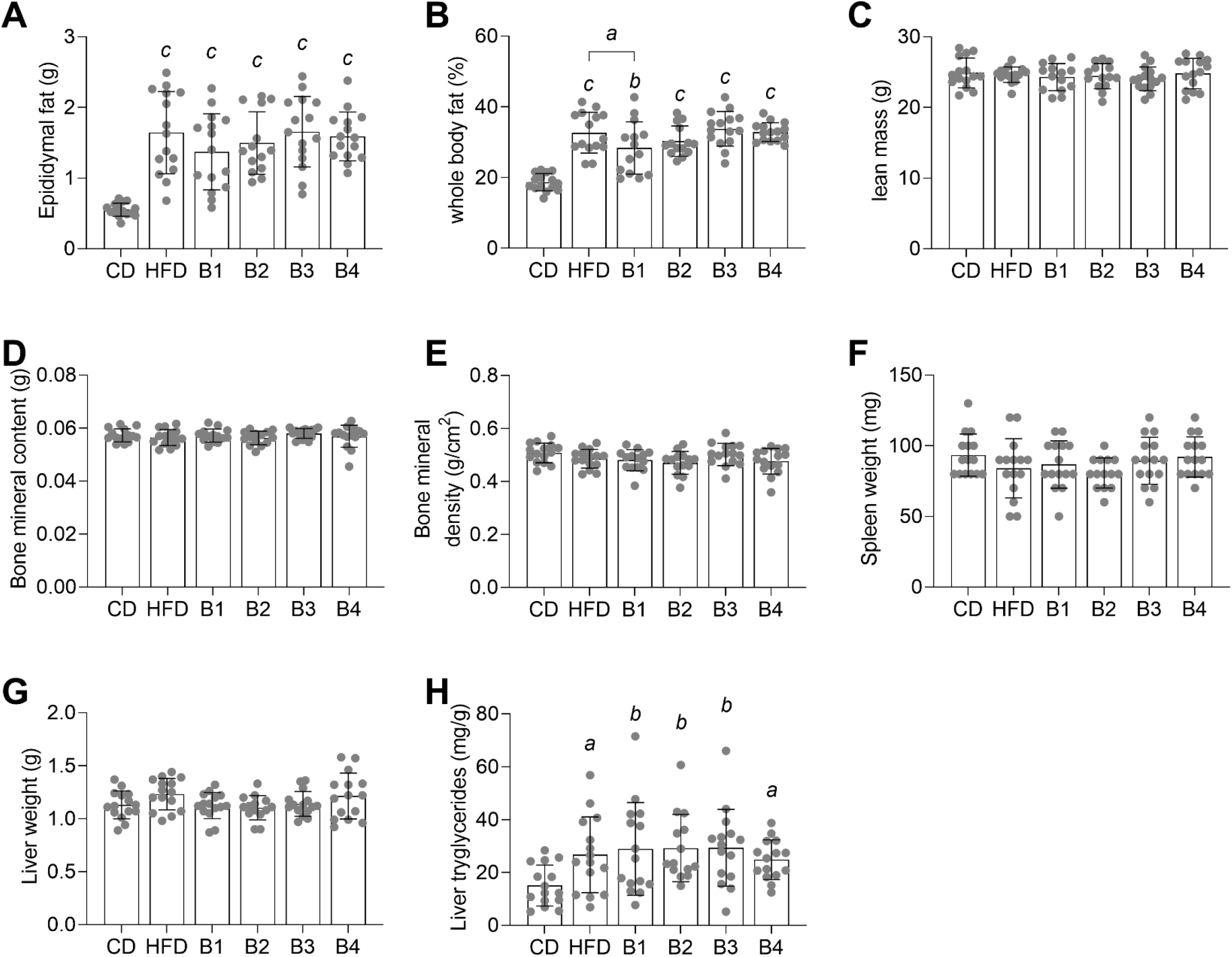
Fat accumulation induced by HFD feeding was reduced by B1 treatment. Gonadal fat surrounding the epididymis (A). Whole body fat (B), lean mass (C), bone mineral content (D) and bone mineral density (E) determined by X-ray absorptiometry. Weights of spleen (F) and liver (G), and triglycerides content in liver (H). Data are shown as mean ± SD, with dots representing each mouse (n=14-15). Letters over bars indicate significant differences (*^a^*P<0.05, *^b^*P<0.01, *^c^*P<0.001) for comparison to CD, based on Fischer’s LSD post-hoc testing following a significant ANOVA effect.

Altogether, these results suggest that treatment with the bacterial strain B1 alleviates the development of metabolic syndrome in HFD-fed mice.

### Behaviour characterization

We then set to determine behavior alterations expected to be induced by HFD (Lizarbe *et al*., 2019; Garcia-Serrano, Vieira *et al*., 2022), and whether treatment with bacteria exerted any protective mechanisms. HFD-fed mice showed unaltered exploration of the elevated plus-maze, a task that evaluates anxiety-like behaviors (figure 4A, supplementary figure S1). However, HFD-fed mice receiving B3 spent less time in closed arms than CD-fed mice (P=0.001 after Kruskal-Wallis P=0.032), suggesting anxiolytic effects of B3.

**Figure 4.**
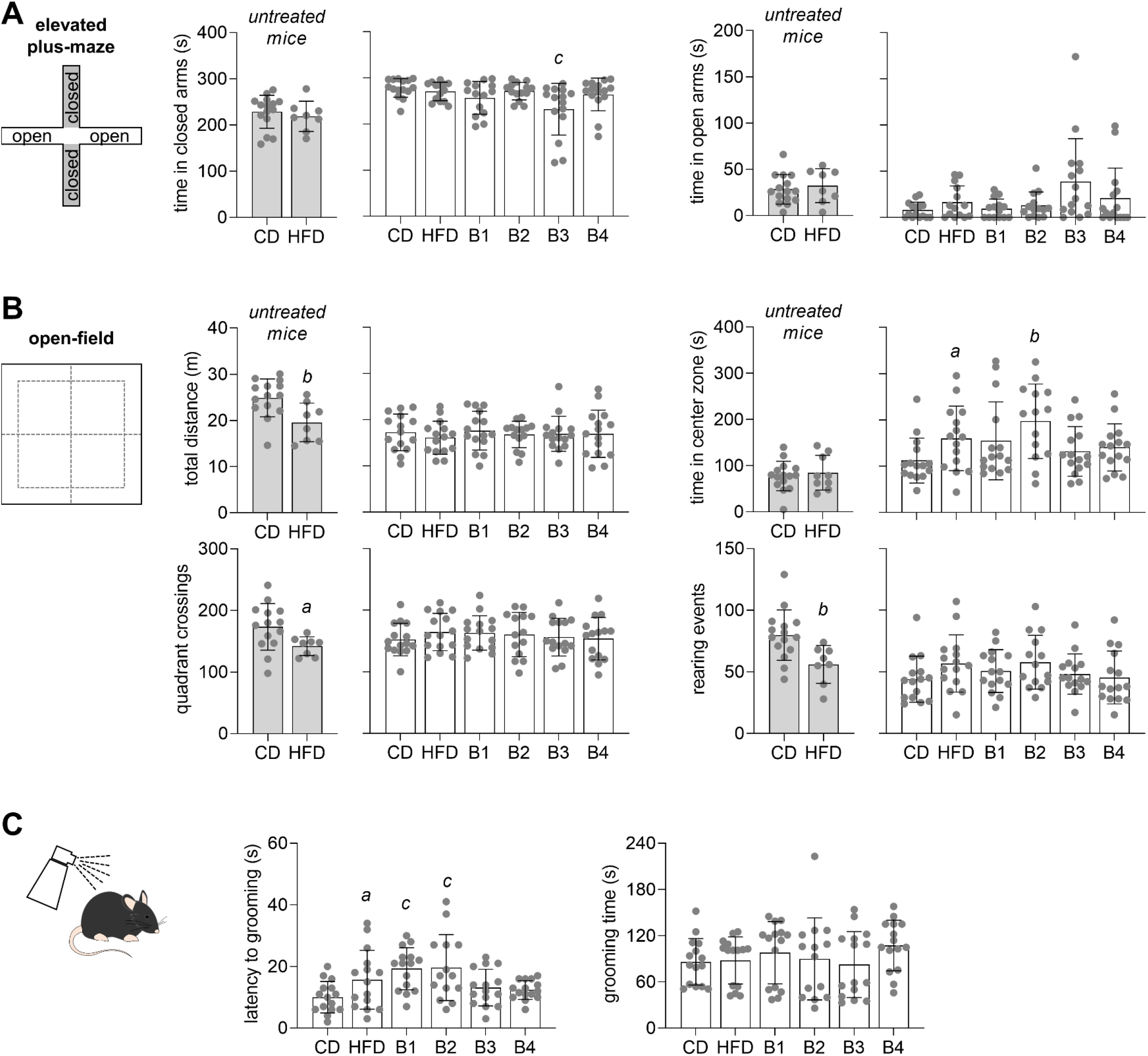
Mouse behavior characterization. (A) Time spent in closed arms, open arms or in center of the elevated plus-maze. (B) Walked distance, number of quadrant crossings, number of rearings, and time spent in center zone during open-filed arena exploration. (C) Latency to grooming and time spent grooming after sucrose spray in the splash tests. Data are mean ± SD, with dots representing each mouse. Letters over bars indicate significant differences (*^a^*P<0.05, *^b^*P<0.01, *^c^*P<0.001) for comparison to CD, based on Student t-tests, or on Fischer’s LSD post-hoc testing following a significant ANOVA effect, or Dunn’s post-hoc testing following significant Kruskal-Wallis test.

We then examined open-field exploration for 8 minutes. As in other models of diet-induced obesity (Garcia-Serrano, Mohr *et al*., 2022), untreated HFD-fed mice showed lower total distance walked in the arena, less quadrant crossings and less rearing events than CD-fed mice, whereas time spent exploring the center of the arena was similar (figure 4B). In contrast, HFD-fed mice that were gavaged with medium or bacteria B2 showed increased exploration of the arena center, as compared to medium-gavaged, CD-fed mice (respectively, P=0.028 and P=0.001 after Kruskal-Wallis P=0.040). Typically, reduced exploration of center *versus* periphery is interpreted as reduced anxiety-like behavior (Prut & Belzung, 2003). Immobility time (freezing episodes) and mouse rotations during open-field exploration were not modified (supplementary figure S2).

We then used the splash sucrose test on bacteria suspension-treated mice to evaluate depressive-like behavior. Although total grooming time after sucrose splash was not altered (Kruskal-Wallis P=0.041), HFD increased the latency to start grooming (ANOVA F=4.18, P=0.002; figure 4C), which was particularly significant for HFD (P=0.036), B1 (P<0.001) and B2 (P<0.001) groups, when compared to controls.

Altogether, these observations indicate that HFD or bacteria supplementations did not cause stress, anxiety or depression-like behaviors.

Finally, we set to test the impact of HFD and bacteria suspension treatments on memory performance. Spatial memory was not impacted by HFD in either untreated (P=0.060) or gavaged mice (ANOVA F=0.331, P=0.89; figure 5A). Exploration of the Y-maze, as depicted by number of arm entries was also similar across experimental groups. Next, we examined novelty recognition memory. After familiarization with 2 objects, mice normally tend to spend more time exploring a novel object, or an object that has been displaced in the arena. When compared to controls, novel object recognition was impaired in mice fed HFD that were untreated (P=0.043). When mice were gavaged with either medium or bacteria HFD did not reduce object recognition (ANOVA F=1.71, P=0.14; figure 5B). Novel location recognition was not impaired by 8 weeks of HFD feeding (*versus* CD) in either untreated (P=0.816) or gavaged mice (ANOVA F=2.21, P=0.061; figure 5C).

**Figure 5.**
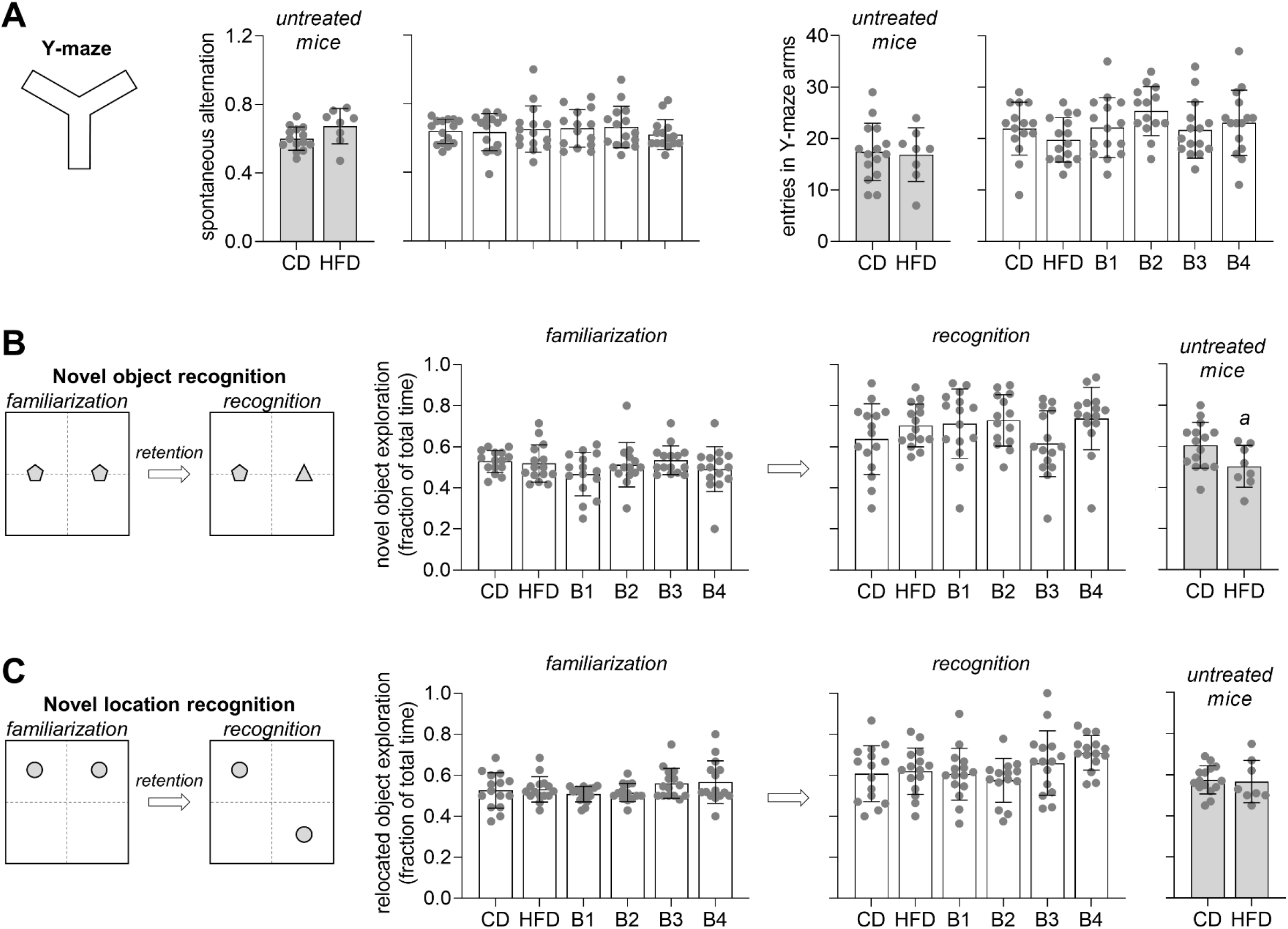
Memory performance in a Y-maze (A) and object recognition (B,C) tests. (A) Spontaneous alternation as fraction of possible alternations in the Y-maze test, and number of entries in the Y-maze arms. (B) Object exploration in novel object recognition during the familiarization (training) and recognition phases (testing). (C) Object exploration in novel location recognition during the familiarization (training) and recognition phases (testing). Object exploration in the familiarization phase was on average ∼50% for each object (graphs on the left). Data are mean ± SD, with dots representing each mouse. *^a^*P<0.05 from a Student t-test comparing HFD and CD.

In sum, despite previous reports on memory impairment in diet-induced obesity models (Lizarbe *et al*., 2019; Garcia-Serrano, Mohr *et al*., 2022; Garcia-Serrano, Vieira *et al*., 2022), mice under treatment with culture medium or bacteria do not show HFD-induced memory impairment in the present study.

### Prediction of bacteria supplementation effects

We used a 5-component PLS regression to discriminate HFD-from CD-fed mice based on all the measured parameters of metabolism and behaviour. The 2 first components were actually able to explain 92% of the total variance in the data, and provided full separation of CD and HFD mice in the score plot (figure 6A), namely along component 1 (figure 6B). VIP scores estimated from the PLS model indicate that fat accumulation and impaired glucoregulation are the most striking effects of HFD feeding among all measured parameters (figure 6C). Coefficients from the PLS model were then used to predict whether individual mice showed a phenotype closer to CD or HFD. Mice treated with B1 showed a larger shift from HFD towards CD than the other 3 bacteria (figure 6A,B).

**Figure 6.**
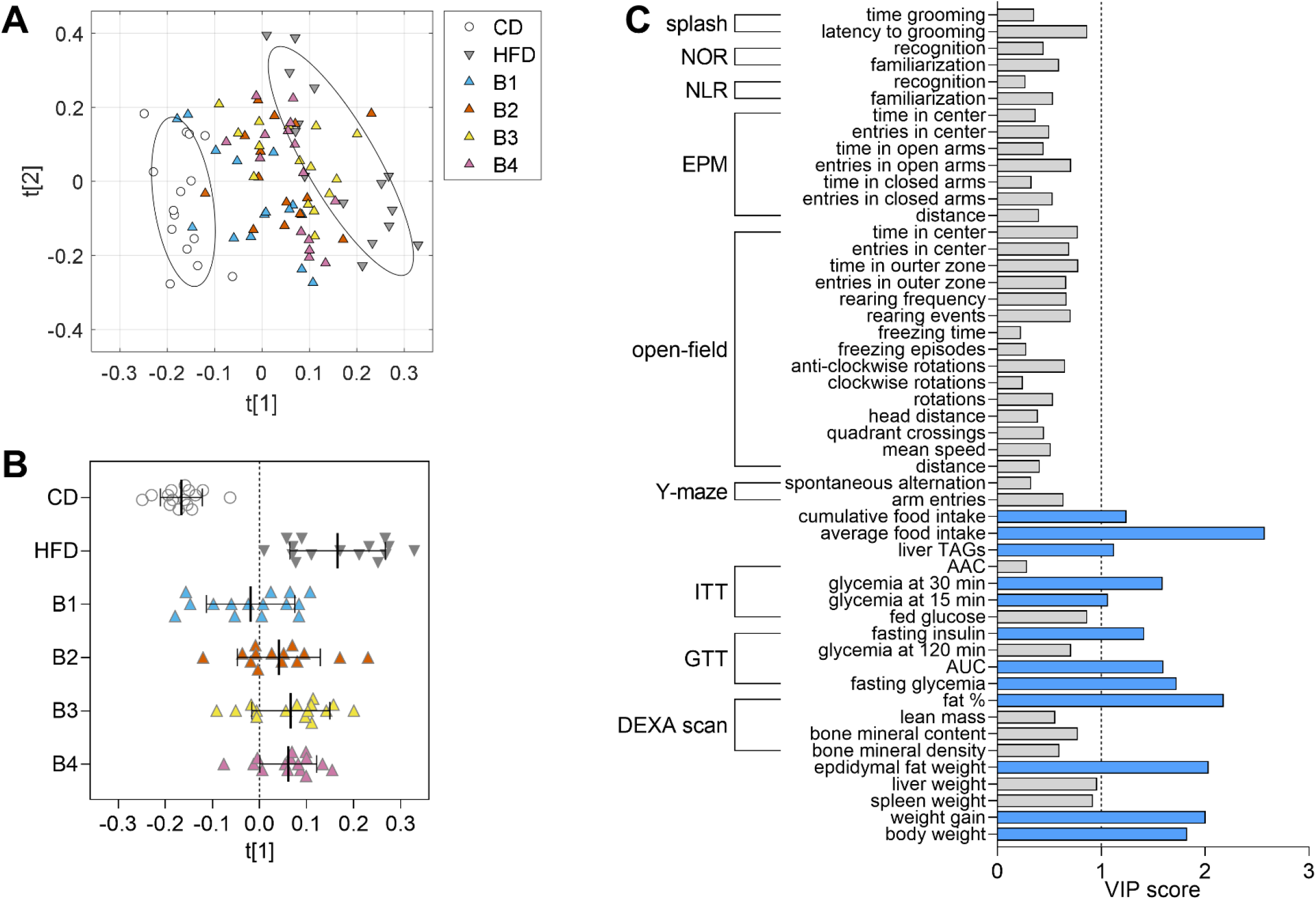
PLS regression of measured parameters to discriminate HFD-fed mice (gray triangles) from controls (open circles). (A) Score plot with each symbol representing one mouse according to components 1 (t[1]) and 2 (t[2]), which explained 83% and 9% of total variance, respectively. Ellipsoids denote group SD for the HFD- and CD-fed mice. The obtained model was used to calculate (predict) the distribution of probiotic-treated mice in the score plot. (B) The score for the first component (t[1]) alone allows to visualize the approximation of B1-treated mice to CD-fed mice. Data are mean ± SD, with symbols representing each mouse (n=14-15). (C) VIP scores calculated from the resulting PLS model for all behaviour and metabolic tests and respective parameters analysed. Blue bars highlight VIP>1.

### Permeability of gut and blood-brain barriers

HFD feeding is proposed to increase permeability of the gut barrier (Müller *et al*., 2016) and BBB (de Paula *et al*., 2020). BBB permeability to a Dextran-based tracer of 70 kDa was assessed by administering the tracer i.p. (figure 7A), and measuring its levels in ventricular blood and in multiple brain areas. Brain-to-blood levels of the tracer varied across brain areas, but were not significantly modified by HFD feeding or bacteria treatments (ANOVA: brain area F=41.32, P<0.001; treatment F=1.60, P=0.201, interaction F=15.66, P=0.249; figure 7B). We tested gut barrier permeability to Dextran-based tracer of 4 kDa administered by oral gavage (figure 7A). Levels of the tracer in plasma collected from the portal vein or by cardiac puncture (ventricular blood) were not significantly modified by HFD feeding or treatments with bacteria (ANOVA: blood source F=12.48, P=0.002; treatment F=1.05, P=0.411, interaction F=1.05, P=0.648; figure 7C).

**Figure 7.**
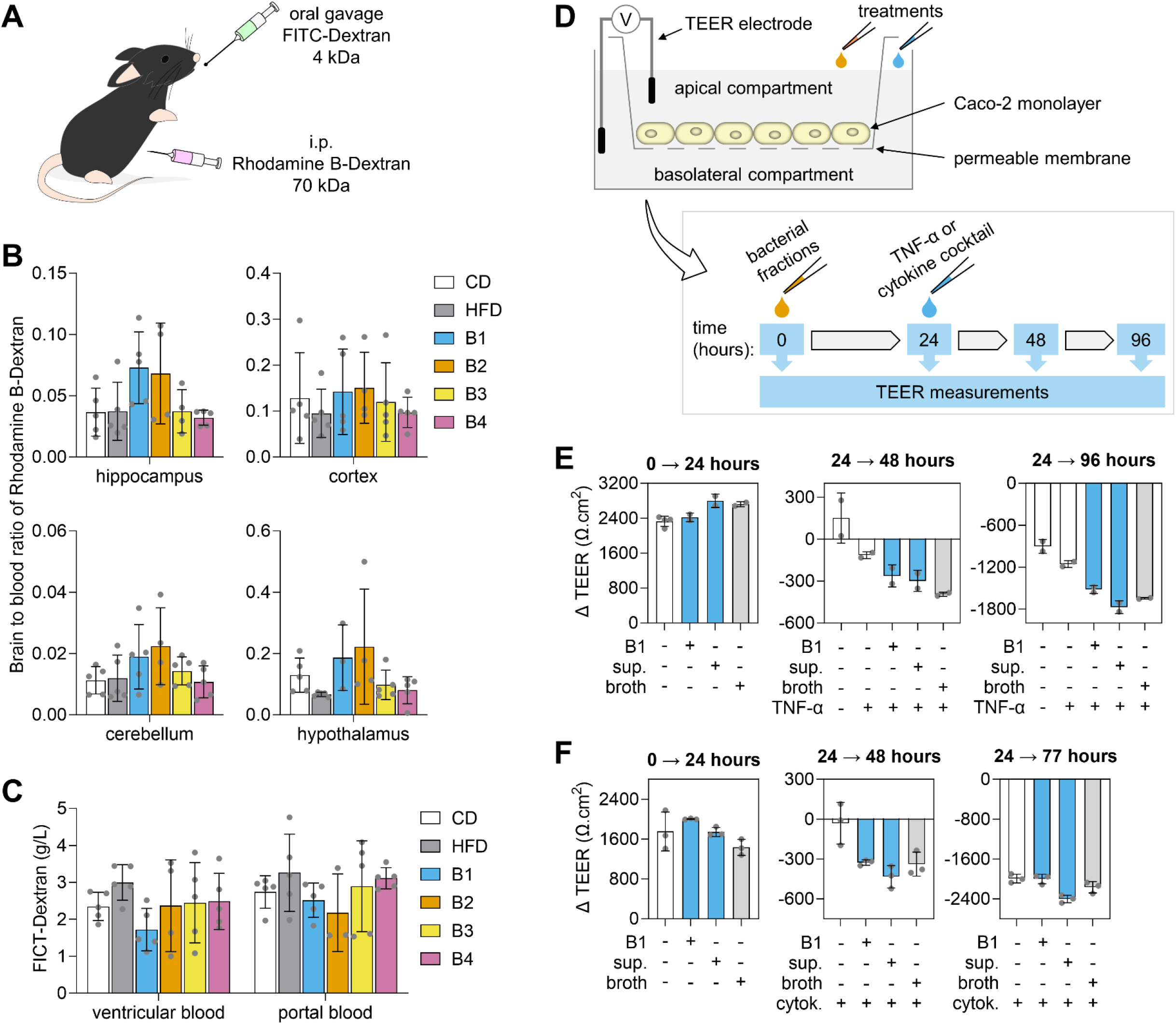
Permeability of gut and blood-brain barriers. (A) Oral and intra-peritoneal (i.p.) administration of dextran tracers for determination of permeability across gut barrier and blood-brain barrier, respectively. (B) Ratio of Rhodamine B-Dextran of 70 kDa levels between brain structures (mg/g wet tissue) and ventricular plasma (g/L). (C) Plasma concentration of FITC-Dextran of 4 kDa in ventricular and portal blood after oral gavage of the tracer, depicting gut permeability. (D) Study design for TEER measurements *in vitro*. (E-F) Change in TEER of Caco-2 cells in response to exposure to heat-inactivated B1, B1 supernatant (sup.) or culture broth for 24 hours, followed by basolateral addition of TNF-α alone (E) or a cocktail of cytokines (cytok.; F). Data are mean±SD, with dots representing each mouse in B-C (n=3-5), or each independent cell culture in E-F (n=2-4).

Given the metabolic health improvements afforded by B1, and the tendency for gut permeability to increase with HFD-feeding, and to be reduced by B1 treatment (figure 7C), we then tested whether B1 was able to increase the transepithelial electrical resistance (TEER) of Caco-2 cells *in vitro* (figure 7D). Due to the anaerobic nature of the bacteria, we were not able to perform experiments with live bacteria. Thus, we tested heat-inactivated bacteria and the supernatant fraction. To be able to assess improvements in cell barrier integrity in response to exposure to bacterial fractions, experiments were started before TEER reached its maximal levels, which in our hands corresponded to around 3000-3500 Ω cm^2^. Exposure of the Caco-2 cells to bacterial fractions for 24 hours did not result in any significant increase in TEER compared to control (figure 7E). Challenging the cell barrier integrity through addition of TNF-α on the basolateral side, decreased TEER with no difference between the bacteria-treated cells and their respective controls. Thus, exposure of polarized Caco-2 cells to these fractions of B1 does not improve the barrier integrity in this experimental setting. A similar result was obtained by adding a cocktail of cytokines, rather than TNF-α alone (figure 7F).

### SCFA production

Fermentation acid profiles of the pure bacterial cultures were tested following growth in the same medium used to pre-grow the bacteria prior to gavage. All strains produced varying concentrations of butyrate and a range of other fermentation acids (figure 8). Strains B1 and B2 both produced acetate while strains B3 and B4 were acetate utilisers, but produced detectable amounts of lactate. B1 produced large amounts of formate. The relatively high amounts of propionate in the basal media were also detected in the culture supernatants (figure 8).

**Figure 8.**
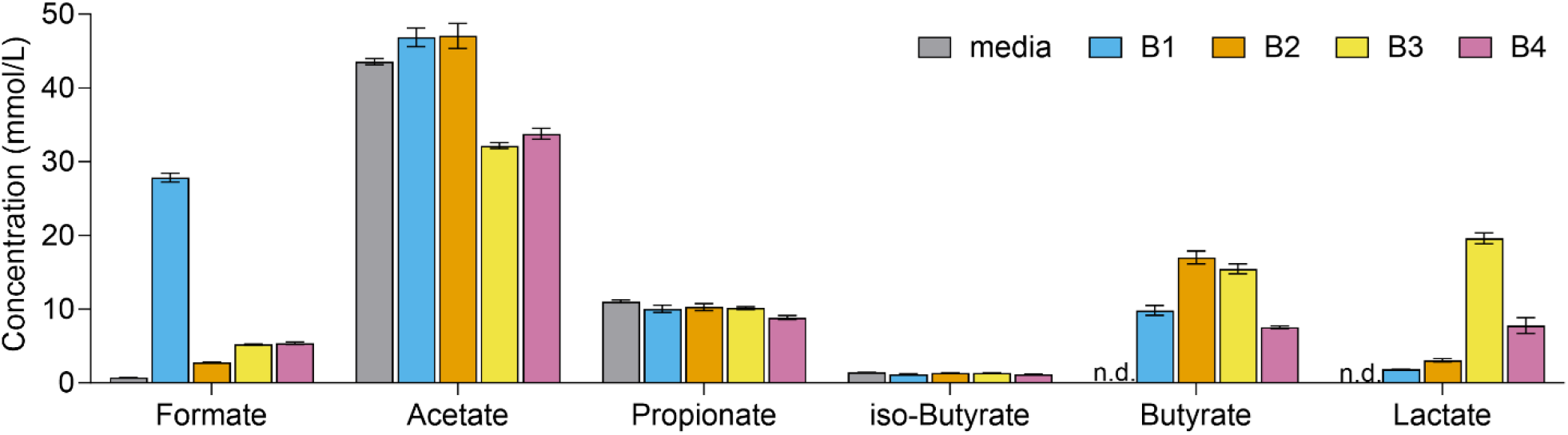
Detection of fermentation acids in culture supernatants following overnight growth. Legend: n.d., not detected.

## Discussion

The present results unveil potential beneficial effects of novel anaerobic bacterial strains on metabolic disease, as well as on modulating of brain function in a model of diet-induced obesity that has impaired glucose tolerance and reduced insulin sensitivity.

Gut microorganisms play an important role in determining obesity development upon feeding a lard-based HFD (Kübeck *et al*., 2016), and several bacterial strains have been tested for improvements in metabolic health, namely ameliorating metabolic syndrome in diet-induced obese mice (*e.g.* Le Roy *et al*., 2022; Huber-Ruano *et al*., 2022). Our findings demonstrate that B1 supplementation has beneficial effects on HFD-induced whole-body fat accumulation, increased fasting glucose, glucose intolerance, reduced glycemic response to insulin. We had postulated that B1 beneficial effects would involve modulation of the gut barrier permeability. Although we have not observed significant alterations of gut permeability, B1-treated mice tended to have a tighter gut barrier, as evidenced by reduced passage of a 4 kDa tracer from the gut into the blood stream. Changes in gut permeability induced by HFD have been proposed to depend on housing conditions (Müller *et al*., 2016), and thus we set to test whether B1 specifically influences permeability of Caco-2 cells in culture. Altogether, our findings from experiments in either mice or Caco-2 cells *in vitro* do not support the notion that gut permeability is modulated by HFD or the probiotics employed. Metabolic effects of B1 on HFD-fed mice might thus involve direct action of B1-released metabolites on target organs. Nevertheless, our finding paves the way for further exploring this bacterial strain for targeting glucose homeostasis and ameliorating of metabolic syndrome, especially in obese individuals.

The second aim of our study was to determine beneficial effects of the butyrate-producing bacteria on brain function. Thus, mice were screened in a battery of behavioural tests. HFD feeding reduced open-field exploratory behaviour in untreated mice, as in other models of diet-induced obesity (Garcia-Serrano, Mohr *et al*., 2022). However, this was not the case in mice that received vehicle (that is, culture medium) or bacteria treatments by oral gavage. In contrast, HFD-fed mice that were treated with medium or the bacteria B2 spent more time than CD-fed mice walking in the unprotected center of the arena, which is consistent with reduced stress or anxiety (Planchez *et al*., 2019). Accordingly, no stress or anxiety-like behaviour in gavaged mice was evidenced by the elevated plus-maze test.

Probiotic-treated mice in the HFD, B1 and B2 groups showed increased latency to start grooming in the splash sucrose test. Although this is often an indication of anhedonia, it was not evidenced by other parameters of depressive-like behaviour (Planchez *et al*., 2019). Namely, B1-treated mice showed neither a reduction in total time spent grooming after the glucose splash, nor increased immobility in the open-field exploration test.

Previous reports from our lab on diet-induced obesity and HFD feeding models (Lizarbe *et al*., 2019; Garcia-Serrano, Mohr *et al*., 2022; Garcia-Serrano, Vieira *et al*., 2022), as well as the present parallel experiments in untreated, CD-/HFD-fed mice indicate that HFD-exposure leads to some degree of memory impairment. However, vehicle-treated HFD-fed mice showed no memory impairment compared to experimentally matched controls. In the absence of memory impairment in any of the 3 tests, we speculate that supplements included in the vehicle (*i.e.* medium used for bacteria culture) provide protection on HFD-induced memory impairment. In fact, the YCFAGSC medium used for culturing the four bacteria is rich in vitamins (Duncan *et al*., 2002) that might promote brain health (Kennedy, 2016), and became a major confounder within our study design. Notably, the culture medium was also rich in the SCFA propionate (see figure 8), which has been recently suggested to exert neuroprotective effects (*e.g.*, Filippone *et al*., 2020; Hou *et al*., 2021; Grüter *et al*., 2023).

Several studies have previously proposed that metabolic disease impacts BBB permeability (see discussion in de Paula *et al*., 2020). In our study, HFD feeding did not cause increased permeability of the BBB. In agreement, de Paula *et al*. have also shown that HFD-induced BBB leakage in the hippocampus occurs early after introducing HFD and is transient (de Paula *et al*., 2020).

In sum, our findings provide the proof-of-concept that B1 can be used to at least prevent or ameliorate the development of obesity and its complications. The mechanism for B1 actions remains to be determined. However, we note that B1 is different from the remaining bacterial strains for its ability to produce substantial amounts of formate in culture (see figure 8). In fact, formate is a host-microbiome metabolite that regulates human physiology, modulates the immune system, and might provide beneficial effects on metabolism (Pietzke *et al*., 2020).

This bacterial strain is a promising candidate for clinical trials, and for developing functional food products or supplements that will have a positive societal impact, that is, they can be a next-generation-probiotics not currently available. Moreover, such probiotics might find a role in precision medicine for modulating gut flora, the microbiome, and for inducing beneficial blood metabolite profiles in specific subsets of diabetes patients.

## Author contributions

Study design: JMND, UA, IM, CH. Experiments and data analysis: AMGS, CS, CH, JMND, ERK, IM, NB, JM, SG. Writing manuscript: JMND, CH. Revising manuscript: CT, IM, KGS, KS, CH. All authors agree with publication of the manuscript.

## Data availability statement

All data from the study are contained within the manuscript.

## Abbreviations

BBB: blood-brain barrier
CD: control diet
DEXA: Dual-energy X-ray absorptiometry
DMEM: Dulbecco’s modified Eagle’s medium
FITC: Fluorescein isothiocyanate
GTT: glucose tolerance test
HFD: high-fat diet
HPA: hypothalamic-pituitary-adrenal
ITT: insulin tolerance test
PBS: phosphate-buffered saline
SCFA: short-chain fatty acid
TEER: transepithelial electrical resistance

## Acknowledgments/grant support

We thank Jenny Rudolfsson for preparation of bacteria suspensions given to mice, and Tina Ovlund for performing the liver TAGs analysis. This work was supported by Sten K Johnsons Stiftelse, Albert Påhlsson Stiftelse and the Swedish Foundation for Strategic Research (SM20-0014 to CH). The Knut and AliceWallenberg foundation, and the Medical Faculty at Lund University are acknowledged for financial support to JMND. The authors acknowledge support from the Lund University Diabetes Centre, which is funded by the Swedish Research Council (Strategic Research Area EXODIAB, grant 2009-1039) and the Swedish Foundation for Strategic Research (grant IRC15-0067). The Strategic Research Area MultiPark (Multidisciplinary Research on Parkinson’s Disease) is ackowledged for access to mouse behaviour labs. The Rowett Institute receives support from the Scottish Government (RESAS).

## Supplementary figures

**Figure S1.**
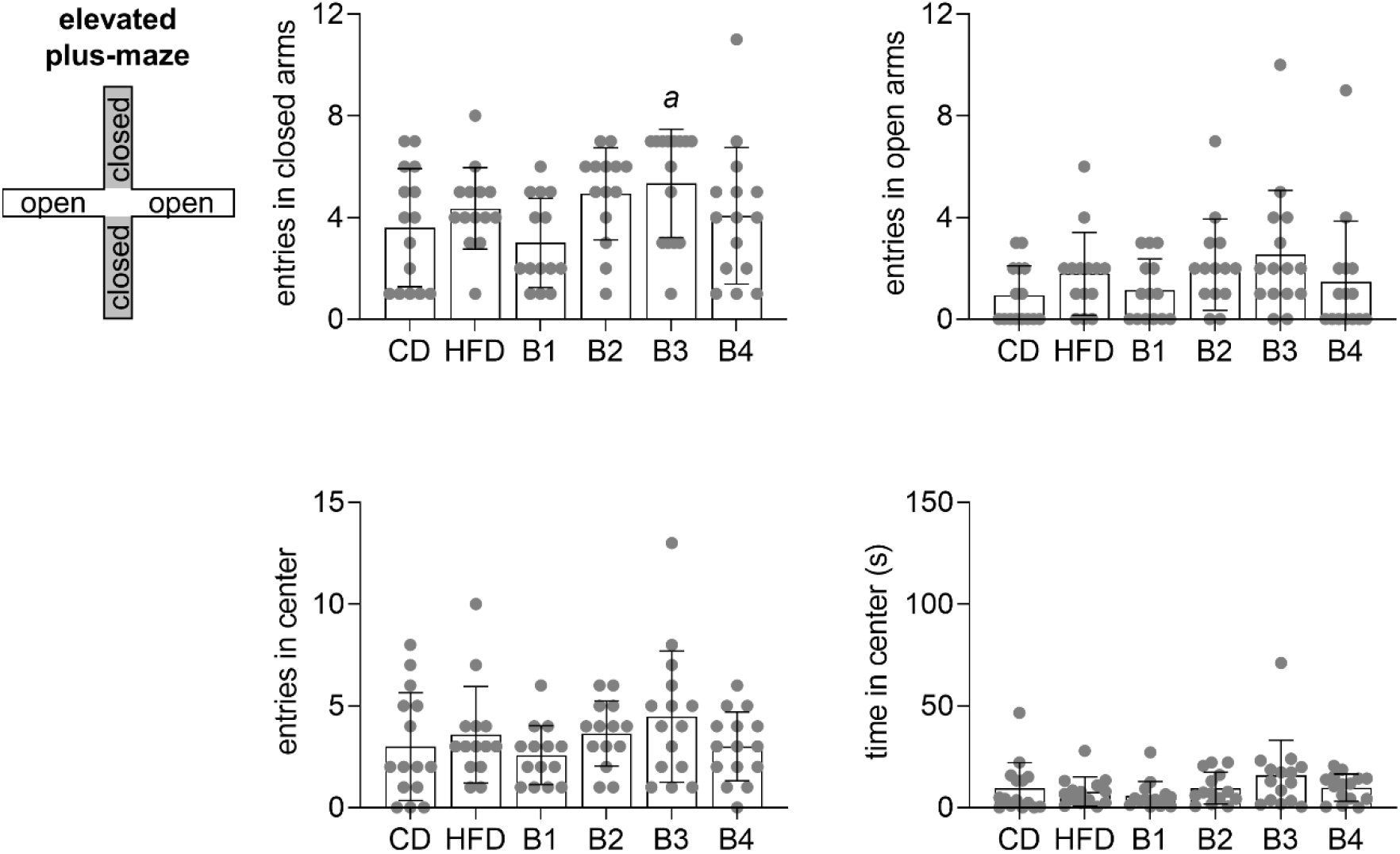
Additional parameters determined during the elevated plus maze test. Data are mean ± SD, with dots representing each mouse. *^a^*P<0.05 for comparison to CD, based Dunn’s post-hoc testing following significant Kruskal-Wallis test.

**Figure S2.**
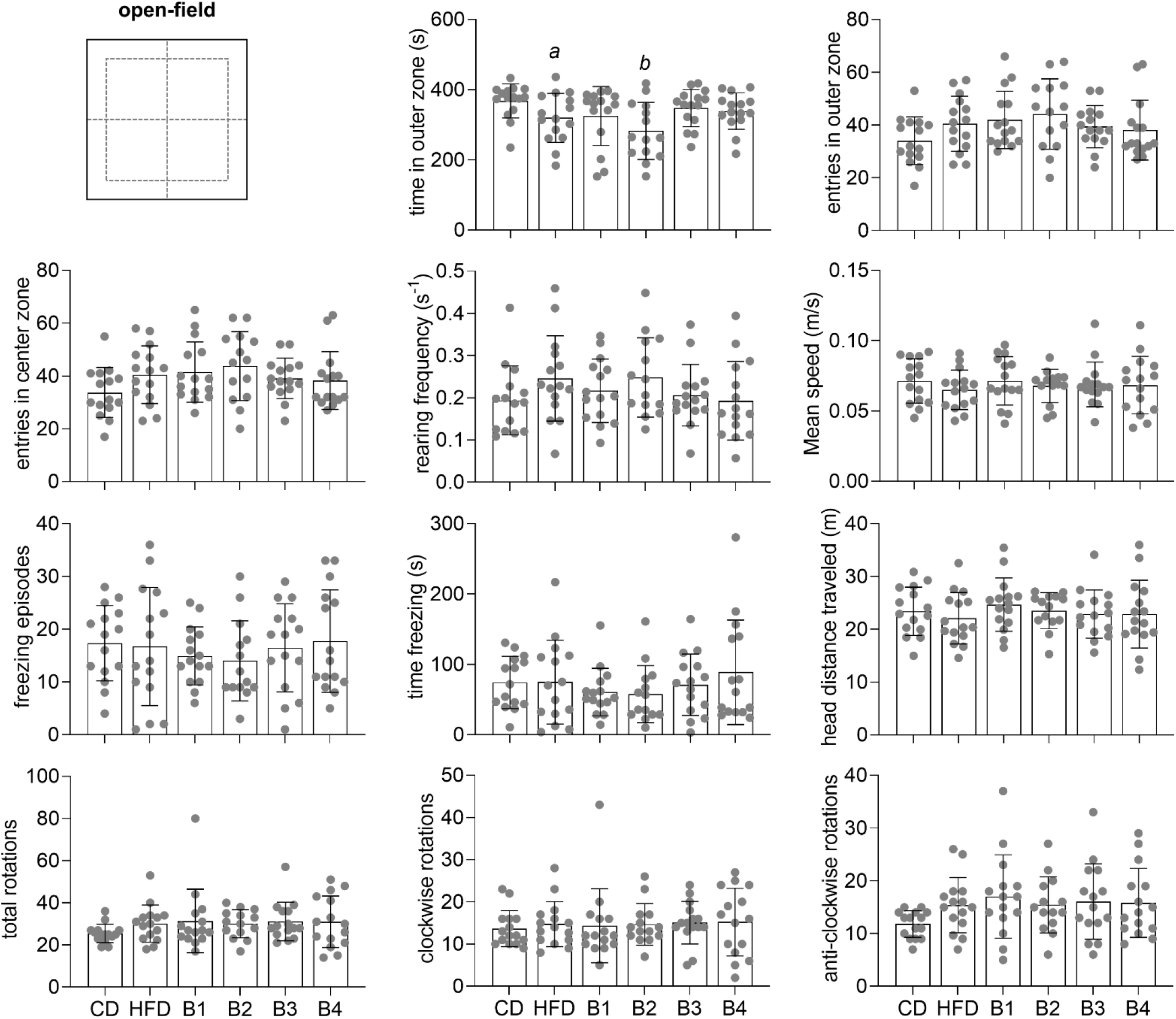
Additional parameters determined during the open-field exploration test. Data are mean ± SD, with dots representing each mouse. Letters over bars indicate significant differences (*^a^*P<0.05, *^b^*P<0.01) for comparison to CD, based on Fischer’s LSD post-hoc testing following a significant ANOVA effect.

